# PCRMS: a database of predicted *cis*-regulatory modules and constituent transcription factor binding sites in genomes

**DOI:** 10.1101/2021.07.23.453290

**Authors:** Pengyu Ni, Zhengchang Su

**Affiliations:** Department of Bioinformatics and Genomics, the University of North Carolina at Charlotte, 9201 University City Boulevard, Charlotte, NC 28223, USA

**Keywords:** Cis-regulatory modules, Database, Transcriptional factor binding sites, Functional analysis

## Abstract

More accurate and more complete predictions of *cis*-regulatory modules (CRMs) and constituent transcriptional factor (TF) binding sites (TFBSs) in genomes can facilitate characterizing functions of regulatory sequences. Here, we developed a database PCRMS (https://cci-bioinfo.uncc.edu) that stores highly accurate and unprecedentedly complete maps of predicted CRMs and TFBSs in the human and mouse genomes. The web interface allows the user to browse CRMs and TFBSs in an organism, find the closest CRMs to a gene, search CRMs around a gene, and find all TFBSs of a TF. PCRMS can be a useful resource for the research community to characterize regulatory genomes.

## Background

*cis*-regulatory modules (CRMs), such as enhancers, promoters, silencers, and insulators, are composed of clusters of short DNA sequences where transcriptional factors (TFs) can bind to regulate the expression of target genes in many biological processes[1]. Recent studies have showed that the vast majority of complex trait-associated single nucleotide variants (SNVs) are located in non-coding sequences (NCSs), and often disrupt transcriptional factor (TF) binding sites (TFBSs) in CRMs[2, 3]. Variation of CRMs also plays a crucial role in divergence in closely related species [4–7]. In principle, variation in TFBSs in a CRM could affect the affinity of cognate TFs, resulting in changes in chromatin modifications and target gene expression in specific cell types in tissues, and ultimately leading to diversity of complex traits, including susceptibility to common complex diseases [8–14]. Therefore, more accurate and more complete categorization of CRMs and constituent TFBSs in the human and important model organisms genomes can greatly facilitate characterizing functions of regulatory sequences and their roles in many important biological processes including disease and evolution.

Recently, a plethora of next-generation sequencing (NGS)-based technologies have been developed to characterize different features of CRMs at a genome scale, such as chromatin immunoprecipitation followed by sequencing (ChIP-seq) [15] to locate TFs binding or histone modifications, as well as DNase I hypersensitive sites sequencing (DNase-seq) [16], assay for transposase-accessible chromatin using sequencing (ATAC-seq)[17], formaldehyde-assisted isolation of regulatory elements sequencing (FAIRE-seq) [18] and micrococcal nuclease digestion with deep sequencing (MNase-seq) [19] to identify the chromatin accessibility. Consequently, an exponentially increase number of datasets have been generated using these technologies by consortia such as ENCODE[20, 21], Epigenomics Roadmap [22, 23] and Genotype-Tissue Expression (GTEx)[24]. Based on different data types that capture different features of CRMs, many computational strategies have been developed to predict CRMs. For instances, based on TF ChIP-seq data, methods such as SpaMo[25], CPModule[26], COPS[27], and INSECT[28] have been developed to identify regions of TF binding as potential CRMs. Based on histone modification marks and chromatin accessibility data, hidden Markov models[29, 30] and dynamics Bayesian models[31] have been developed to predict CRMs and their functional states in different cell types. Based on bidirectionally transcribed pairs of capped RNAs, or enhancer RNA (eRNA), the FANTOM 5 project identified active enhancers in various human and mouse tissues [32]. By integrating multiple tracks of epigenetics marks, TF binding, and predicted and experimentally validated enhancers, several groups have developed CRM/enhancer databases, such as dbSUPER[33], SEdb[34], DENdb[35], EPDnew promoters[36], UCNEbase[37], CraniofacialAtlas[38], GeneHancer[39], HACER[40], RAEdb[41], HEDD[42], DiseaseEnhancer [43], SEA[44], EnhancerAtlas [45], and SCREEN[46]. However, these databases only cover a small portion of enhancers/CRMs encoded in the genomes, and some may have a high false positive rate [47]. For instance, even the most currently updated SCREEN database that stores candidate *cis*-regulatory elements (cCREs) predicted by the ENCODE phase 3 consortium contains only 926,535 and 339,815 cCREs in the human and mouse genomes, with a mean length of 273bp and 272bp, respectively [46], which is much shorter than the mean length (~2,000bp) of known human and mouse CRMs in the VISTA database[48], indicating that cCREs in both genomes might be underpredicted. Moreover, none of these databases provide *de novo* predicted TFBSs in enhancers/CRMs, which are critical to understand the mechanisms of transcriptional regulation, and to pinpoint causal variants of phenotype diversity and disease risks.

Using a highly efficient CRM and TFBS prediction tool dePCRM2 that we developed recently [47], we have predicted CRMs and constituent TFBSs in human (*Homo sapiens*) and mouse (*Mus musculus*) using a large number of TF ChIP-seq datasets in the organisms. Comparative analysis indicates that our predictions are substantially more accurate and more complete than those in existing databases [47]. To facilitate the research community to use these predictions for various purposes, we constructed an online database PCRMS. The database currently contains 1,404,973 and 920,068 CRM candidates (CRMCs) and 90,671,016 and 104,251,155 TFBSs for 201 and 210 unique motif (UM) families in human and mouse, respectively. To our best knowledge, these represent the most complete collections of accurately predicted CRMs and constituent TFBSs in human and mouse genomes. The web interface to PCRMS allows a quick search, browse and visualization of the contents of the database, and provides three functional analysis modules. Using these modules, the users can find the closest CRMs to a gene; search CRMs that are located in a specified region around a gene; and search TFBSs in CRMs for a given TF. The interface also provides copy, export and download functions of selected CRMs or all the predicted CRMs in a BED format. PCRMS will facilitate the research community’s efforts to characterize the regulatory genomes in important organisms.

## Construction and content

### Datasets

We downloaded (10/25/2018) 6,092 TF ChIP-seq datasets for 779 TFs in 468 cells/tissues/organs of humans, and 4,786 TF ChIP-seq datasets for 501 TFs in 162 cells/tissues/organs of mice from the CISTROME database[49]. After filtering out peaks with low quality, for each left peak, we extracted 1,000bp genome sequence centering on the middle of the binding peaks, thereby extending majority of the binding peaks as most called binding peaks have a length short than 1,000bp [47].

### Prediction of CRMs and constituent TFBSs

To predict CRMs and TFBSs, we applied our dePCRM2 algorithm[47] to the datasets with extended binding peaks from each organism using the default parameters. Briefly, dePCRM2 first finds overrepresented motifs and co-occurring motifs pairs (CPs) in each dataset in an organism. It then identifies unique motifs (UMs) by combining highly similar motifs in CPs across all the datasets in the organism. To model the interactions among cooperative TFs, dePCRM2 constructs an interaction network, where UMs are the nodes, and two nodes are connected by a weighted edge with the weight being their interaction score, defined as,

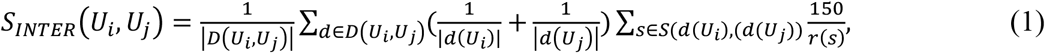

where *D*(*U*_*i*_, *U*_*j*_) is the datasets in which TFBSs of both *U*_*i*_ and *U*_*j*_ occur, *d*(*U*_*k*_) the subset of dataset *d*, containing at least one TFBS of *U*_*k*_, *S*(*d*(*U*_*i*_), (*d*(*U*_*j*_)) the subset of *d* containing TFBSs of both *U*_*i*_ and *U*_*j*_, and *r*(*s*) the shortest distance between any TFBS of *U*_*i*_ and any TFBS of *U*_*j*_ in a sequence *s*. Clearly, *S*_INTER_ allows flexible adjacency and orientation of TFBSs in a CRM and at the same time, it rewards motifs with binding sites co-occurring frequently in a shorter distance in a CRM. Next, dePCRM2 connects two adjacent TFBSs of the UMs if their distance *d* ≤ 300bp, and predicts each resulting connected DNA segment as a CRM candidate (CRMC), thereby partitions the genome regions covered the extended peaks in a CRMC set and a non-CRMC set. Finally, dePCRM2 evaluates each CRMC containing *b*_1_, *b*_2_ …, *b*_*n*_ TFBSs by computing a score defined as,

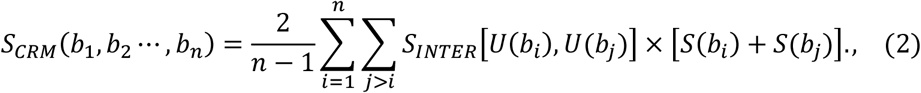

where *U*(*b*_*k*_) is the UM of TFBS *b*_*k*_, *W*[*U*(*b*_*i*_), *U*(*b*_*j*_)] the interaction score between *U*(*b*_*i*_) and *U*(*b*_*j*_), *s*(*b*_*k*_) the binding score of *b*_*k*_ based on the position weight matrix (PWM) of *U*(*b*_*k*_). Only TFBSs with a positive score are considered. dePCRM2 also computes a p-value for each CRMC as follows. For each predicted CRMC, dePCRM2 generates a Null CRMC that has the same length and 4-mer nucleotide frequencies as the CRMC using a third order Markov chain model [50], and computes a S_CRM_ score for each Null CRMC based on a random interaction network which is generated by randomly rewiring the nodes of the UM interaction network. Then, an empirical p-value for a CRMC with a *s*_*CRM*_=s is computed based on the distribution of S_CRM_ score of the Null CRMCs,

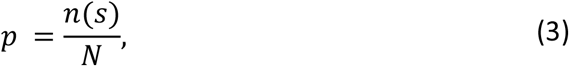

where *n*(*s*) is the number of Null CRMCs with a *s*_*CRM*_ score greater than s, and N total number of the CRMCs.

### Technical implementation

The current version of PCRMS (v2) was developed using MySQL 5.7.17 (http://www.mysql.com) and it runs on a Linux-based Apache2 server (http://www.apache.org). The PHP 7.4 (http://www.php.net) was used for back-end scripting. The interactive interface and responsive features were implemented using Bootstrap 4 (https://getbootstrap.com/), JQuery (http://jquery.com) and dataTables (https://datatables.net/). NCBI sequence viewer 3.38.0 (https://www.ncbi.nlm.nih.gov/projects/sviewer/) was used for visualization.

## Utility and discussion

### Predicted CRMs and constituent TFBSs in the human and mouse genomes

Applying dePCRM2 to the TF ChIP-seq datasets available to us (6/1/2019) in each organism, we predicted 1,404,973 and 920,068 CRMCs in the human[47] and mouse genomes, respectively, comprising 44.03% and 50.39% of their respective genomes. These CRMCs contain 90,671,016, and 104,251,155 TFBSs, comprising 16.71 % and 20.34% of the human and mouse genomes, respectively. As shown in Table 1, our predicted numbers of CRMCs are much larger than those predicted by the ENCODE[46] or GeneHancer[39] teams, our predicted CRMCs also comprise much larger proportions of the genomes than do those predicted by the two teams. Moreover, our predicted CRMCs contain information of *de novo* predicted TFBSs, which are lacking in those predicted by the two teams.

**Table 1.**
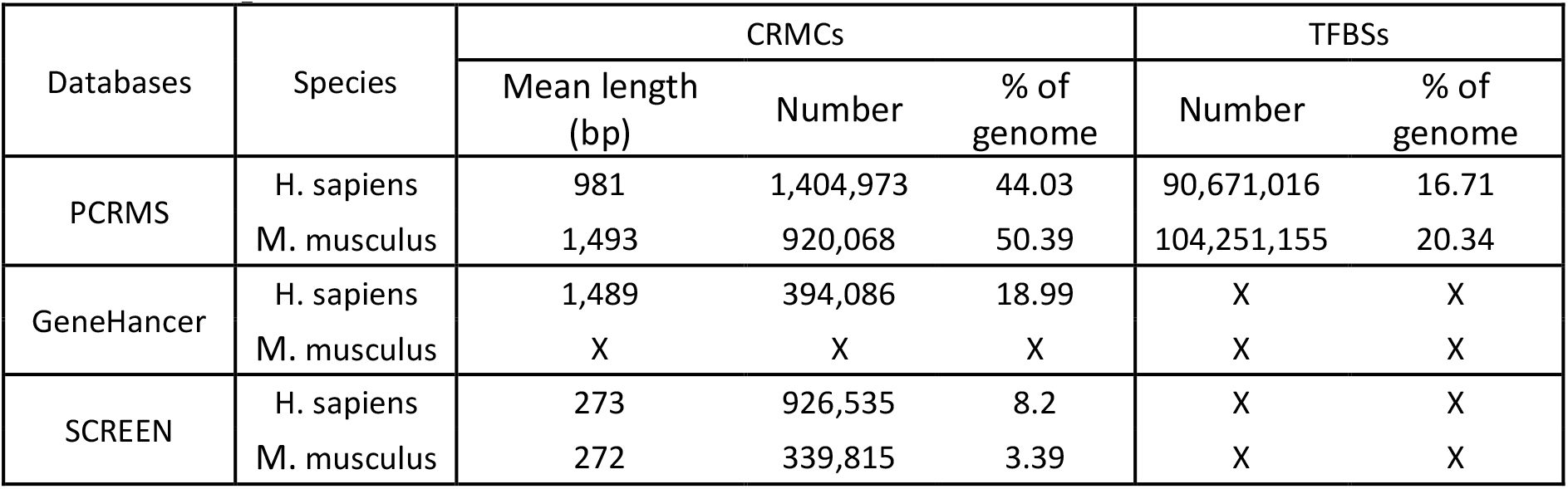
Comparison of PCRMS with the other databases in contents.

dePCRM2 computes a p-value for each predicted CRMCs, and we have shown earlier that the smaller the p-value, the longer the CRMC and the stronger evolutionary constraint the CRMC is subject to [47]. To assist the users who might be interested in CRMCs with different statistical significance, or with different evolutionary constraints, we provide four options of p-value cutoffs (p-value <0.05, 0.01, 5 × 10^−6^ and 1 × 10^−6^) to query the database. Table 2 summarizes the predicted CRMs using these p-value cutoffs, they are subsets of the CRMCs with different length distributions and conservation levels [47]. We have shown that the predicted CRMCs and TFBSs in the human genome are highly accurate based on validations using multiple independent data [47], and the same is true for the predicted CRMCs and TFBSs in the mouse genome (manuscript in preparation, P.N. and Z.S).

**Table 2.** Summary of the predicted CRMs at different p-values in the human and mouse genomes.

| Species | p-value | CRMs |  |  | TFBSs |  |
| --- | --- | --- | --- | --- | --- | --- |
|  |  | Mean length (bp) | Number | % of genome | Number | % of genome |
| H. sapiens | 0.05 | 1,162 | 1,155,151 | 43.47 | 89,948,206 | 16.54 |
|  | 0.01 | 1,292 | 1,020,679 | 42.72 | 88,912,654 | 16.32 |
|  | 5.00E-06 | 2,292 | 428,628 | 31.81 | 71,478,114 | 12.88 |
|  | 1.00E-06 | 2,624 | 327,396 | 27.82 | 64,136,635 | 11.47 |
| M. musculus | 0.05 | 1,749 | 777,409 | 49.9 | 103,718,473 | 20.21 |
|  | 0.01 | 1,944 | 688,033 | 49.06 | 102,730,265 | 19.99 |
|  | 5.00E-06 | 3,182 | 338,635 | 39.53 | 88,579,892 | 16.96 |
|  | 1.00E-06 | 3,780 | 250,606 | 34.75 | 80,002,349 | 15.2 |

### Web interface to the database

We provide a user-friendly web interface to the PCRMS database for quickly inquiring and browsing predicted CRMs and TFBSs at different statistically significant levels in each organism as well as three functional analysis modules. Using these modules, the user can (i) search the closest CRM to a given gene; (ii) search all CRMs in upstream and/or downstream regions of a gene of interest; and (iii) search the TFBSs of a TF on one or more chromosomes (Figure 1).

**Figure 1.**
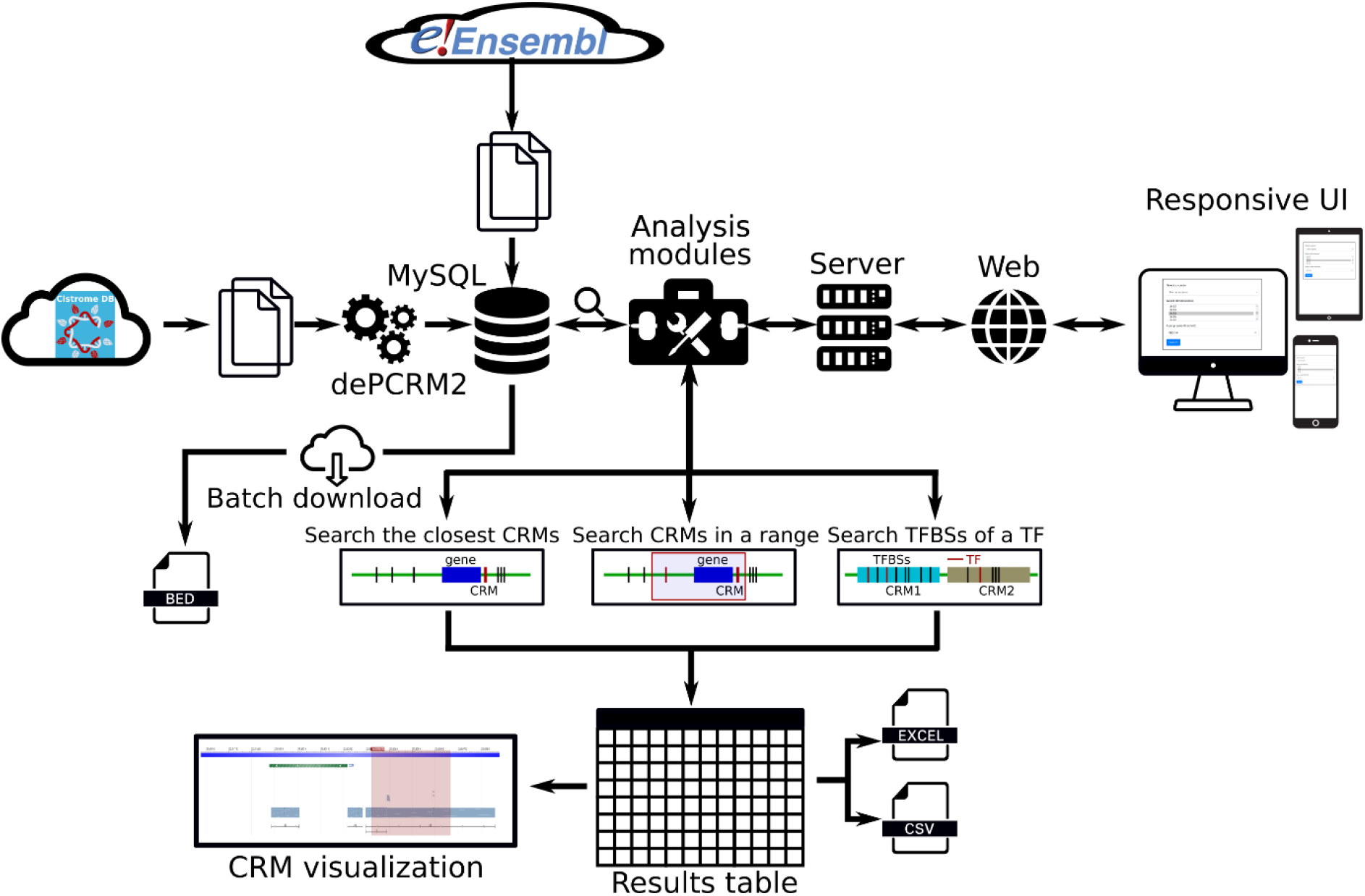
Overview of data integration, analysis modules and features of the PCRMS database.

### Browse of database contents

We provide a Browse function by which the user can browse all CRMs predicted at a selected p-value cutoff on one or multiple selected chromosomes in a selected organism, and inspect each CRMs and constituent TFBSs in detail. The user starts in the search form (Figure 2A) by selecting an organism (e.g., *H. sapiens*), one or more chromosomes (e.g., chrX), and a p-value cutoff (e.g., 1E-06). The search returns all the predicted CRMs (n=8,762) on the chromosome (chrX) of the organism (*H. sapiens*) in the interactive CRM list table (Figure 2B). Clicking on a CRM of interest, e.g. the first CRM hse1000017 in the table pops up the CRM information table (Figure 2C), where some parameter of the CRM are shown in the left panel and the locus of the CRM is displayed in the NCBI sequence viewer (shadowed rectangle) in the right panel, enabling detailed inspections of the genomic context of the CRM, including its neighboring genes and other annotations using the zooming and the translation functions of the viewer. For instance, the viewer reveals that hse1000017 is located in the 2^nd^ through the 5^th^ introns, and spans the 3^rd^ through 5^th^ exons, of the *BCOR* gene that codes for a corepressor of a transcription repressor BCL6, both genes are involved in B lymphocytes differentiation[51, 52]. Interestingly, hse1000017 overlap two regulatory sequences annotated as “enhancer” and “transcriptional *cis*-regulation”, and many ClinVar variants are located in hse1000017 (Figure 2C). Finally, clicking on the CRM ID (e.g., hse1000017) in the right panel of the CRM information table(Figure 2C) will display the CRM’s 5,094 constituent TFBSs in the interactive TFBS table (Figure 2D), which includes the coordinates of the TFBSs, their Umotif IDs, binding scores, Umotif logos and matched known motifs. The vast majority of these TFBSs match those of known TF families (Figure 2D).

**Figure 2.**
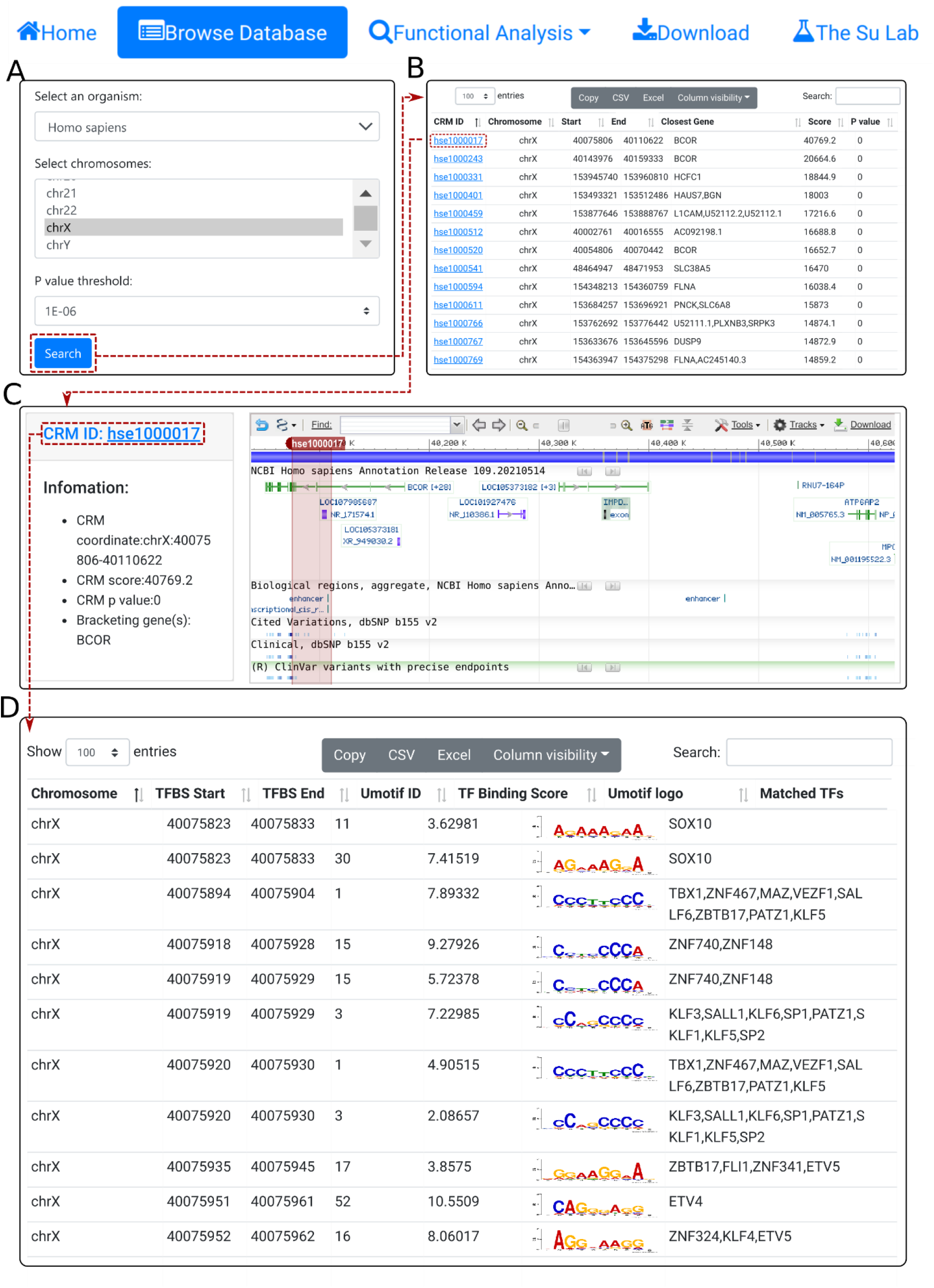
The browse function. **A.** In the search form, the use selects an organism (e.g., *H. sapiens*), one or more chromosomes (e.g., chrX), and a p-value cutoff(e.g., 1E-06). **B.** The searching results are displayed in the CRM list table. Shown is a snapshot of the resulting CRM list table containing 8,762 predicted CRMs on chrX of *H. sapiens*. The first CRM hse1000017 in the list table is selected for further visualization. **C.** In the CRM information table, some parameters of the selected CRM hse1000017 is shown in the right panel, and the locus is displayed in the NCBI sequence viewer for further inspection. Clicking on “hse1000017” in the right panel of the CRM information table displays its constituent TFBSs. **D.** A snapshot of the TFBS table of hse1000017 containing its 5,094 constituent TFBSs.

In both the interactive CRM list table(Figure 2B) and the interactive TFBS table (Figure2C), the user can change the number of entries to display in a page, sort results based on different columns, filter the results using the search box, and set visible columns. The user can copy or export the selected items in a file in the CSV or Excel formats, or export all records if no item is selected by default (Figure 2).

### Functional analyses

To facilitate analyzing potential CRM-gene relationships and TFBSs landscape of specific TFs, we provide three functional analysis modules. First, using the “select closest CRMs to a gene” function, the user can search the closest CRMs to a gene (e.g., *GL13*) in an organism (e.g., *H. sapiens*) at a p-value cutoff (e. g., 1 × 10^−6^)(Figure 2C). The search returns the interactive CRM list table containing all CRMs to which the gene is the closest among all other genes in the chromosome (Figure 2B). In the example of the *GLI3* gene, a total of 53 CRMs are returned. The user can inspect any of them by clicking on the CRM ID, which pops up the information table of the CRM as we demonstrated earlier (Figure 2C). In instance, clicking on the third CRM hse1003435 in the table displays it in the NCBI sequence viewer, revealing that the CRM is located in the 3^rd^ and 4^th^ introns, and spans the 4^th^ and 5^th^ exons, of the *GL13* gene (shadowed rectangle in Figure 3C). Interestingly, hse1003435 overlaps two annotated enhancers and many ClinVar variants (Figure 3C). Finally, clicking on the CRM ID hse1003435 in the right panel of the CRM information table(Figure 3C) displays the CRM’s 988 constituent TFBSs in the interactive TFBS table (Figure 3D). Most of these TFBSs match those of known TF families, while a few need to be determined (TBD) for their cognate TFs(Figure 3D).

**Figure 3.**
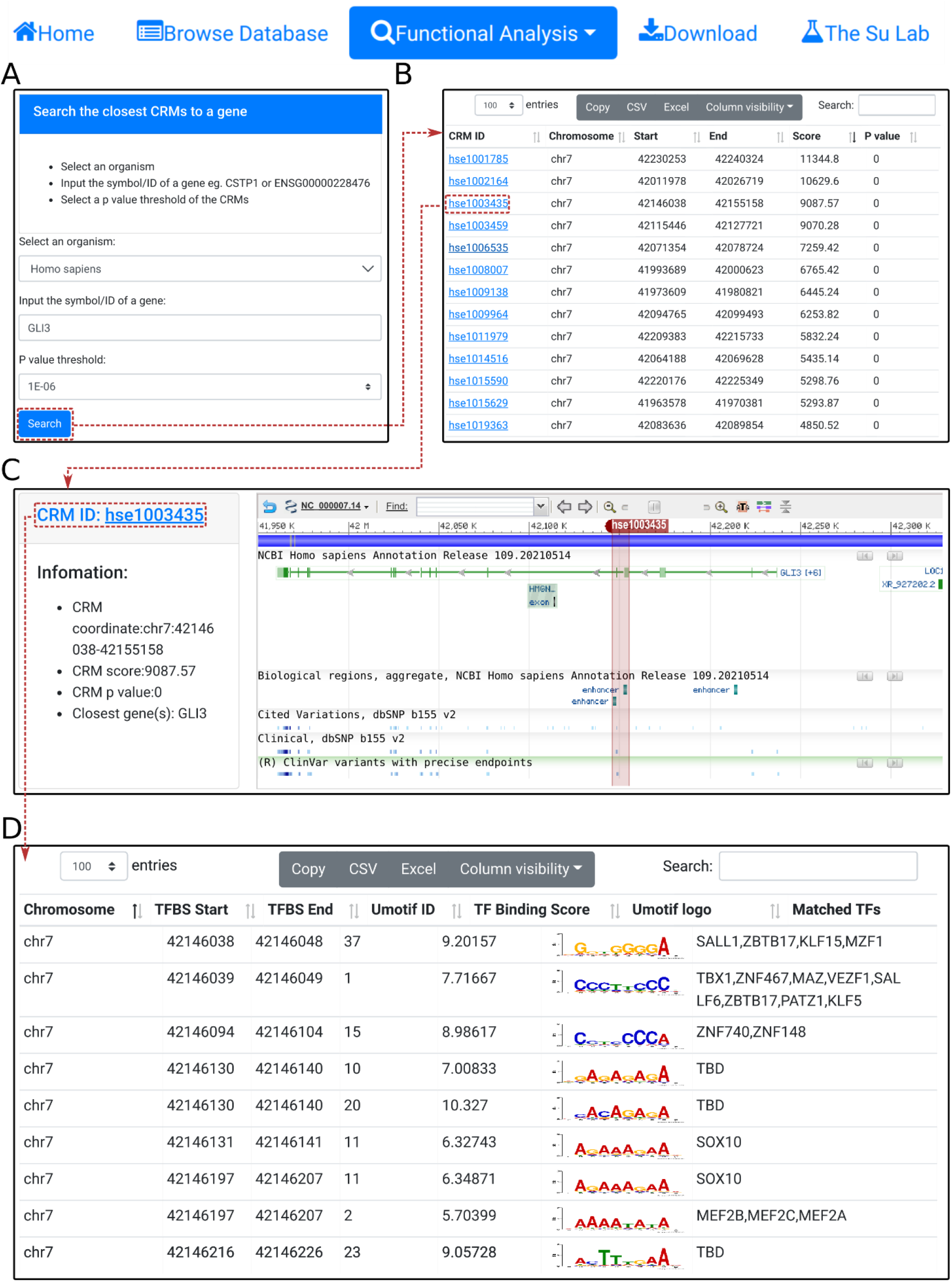
Search the closest CRM(s) to a gene. A. In the search form, the use selects an organism (e.g., *H. sapiens*) and a p-value cutoff (e.g., 1E-06), and inputs a gene name (e.g., *GL13*). **B.** The searching results are displayed in the CRM list table. Shown is a snapshot of the returned CRM list table containing 53 predicted CRMs. The third CRM hse1003435 in the list table is selected for further inspection. **C.** In the CRM information table, some parameters of the selected CRM hse10003435 is displayed in the right panel, and the locus is displayed in the NCBI sequence viewer for further inspections. Clicking on “hse1003435” in the right panel of the CRM information table displays its constituent TFBSs. **D.** A snapshot of the TFBS table of hse1003435 containing its 988 constituent TFBSs.

Second, using the “select CRMs around a gene” function, the user can search in an organism (e.g. *H. sapiens*) all CRMs in upstream and/or downstream regions (e.g., 0.5Mbp) of a given gene (SOX2)(Figure 4A). The search returns all CRMs in the interactive CRM list table (Figure 4B). As before, each CRM can be inspected in its information table by clicking on the CRM ID. In the example of the *SOX2* gene of *H. sapiens*, a total of 102 CRMs on chr3 are returned in the CRM list table(Figure 4B). Inspection of the second CRM hse1002109 in the NCBI sequence viewer reveals that the CRM is located in the 6^th^ and 7^th^ intron, as span the 7^th^ exon, of the *SOX2* gene. Interestingly, it overlaps two annotated enhancer sequences, as well as many ClinVar variants (Figure 4C). Clicking on the CRM ID hse1002109 in the right panel of the CRM information table (Figure 4C) displays the CRM’s 1,344 constituent TFBSs in the interactive TFBS table (Figure 4D). Some of these TFBSs match those of known TF families, while others need to be determined (TBD) for their cognate TFs.

**Figure 4.**
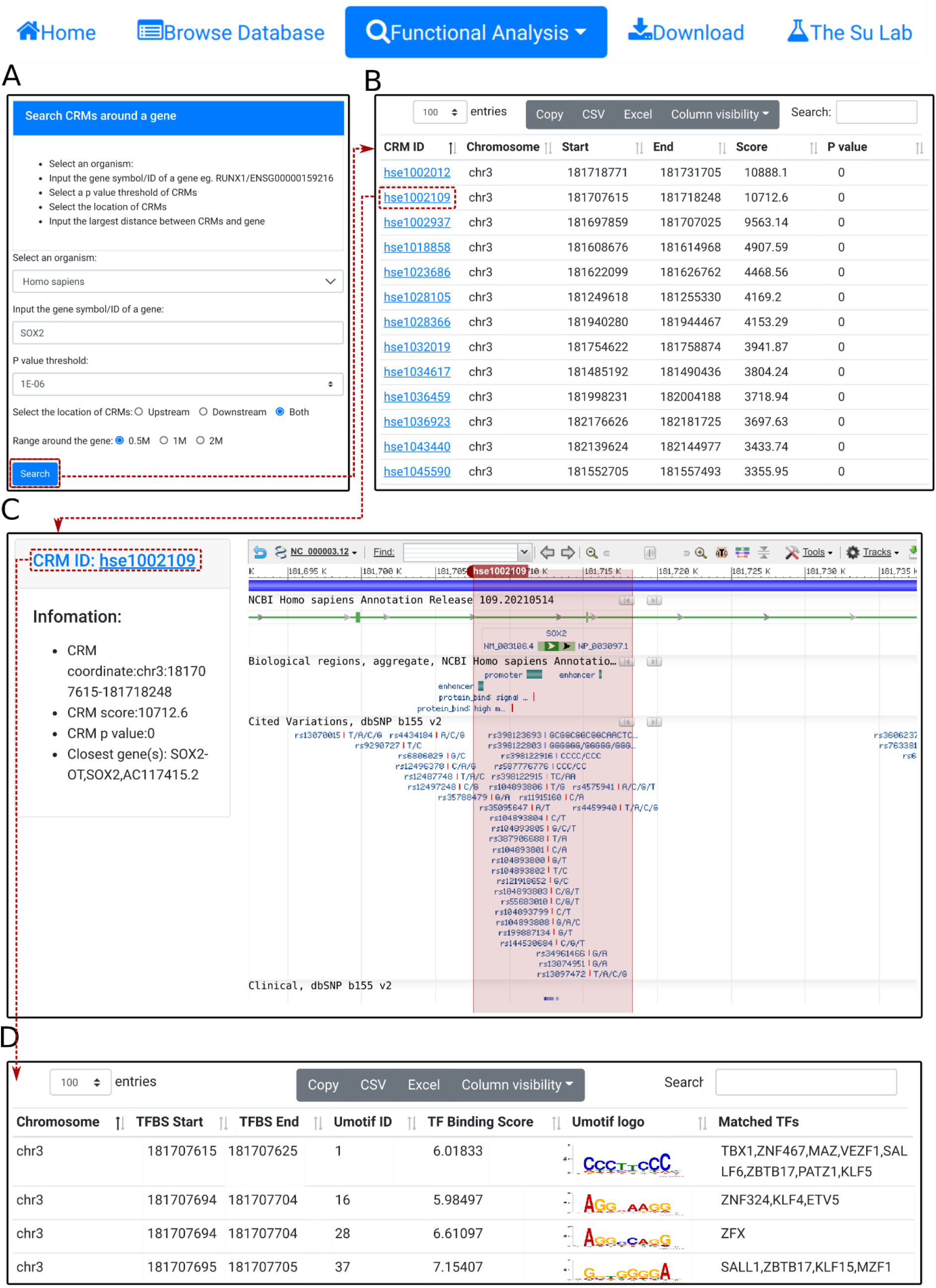
Search CRM(s) in a region around a gene. A. In the search form, the use selects an organism (e.g., *H. sapiens*) and a p-value cutoff(e.g., 1E-06), and inputs a gene name (e.g.,SOX2). **B.** The searching results are displayed in the CRM list table. Shown is a snapshot of the 102 returned CRMs in the table. The second CRM hse1002109 in the list table is selected for further inspection. **C.** In the CRM information table, parameters of the selected CRM hse1002109 is shown in the right panel, and the locus is displayed in the NCBI sequence viewer for further inspections. Clicking on “hse1002109” in the right panel of the CRM information table displays its constituent TFBSs. **D.** A snapshot of the TFBS table of hse1002109 containing its 1,344 constituent TFBSs.

Using the “search TFBSs of a transcription factor” function, the user can retrieve all TFBSs of a given TF (e.g. RUNX1) in one or more selected chromosomes (e.g., chrX) in an organism (e.g., *H. sapiens*) (Figure 5A). The results are returned in the interactive TFBS table (Figure 5B). In the example of the TF RUNX1, a total of 2,678 binding sites are found in chrX of *H. sapiens*.

**Figure 5.**
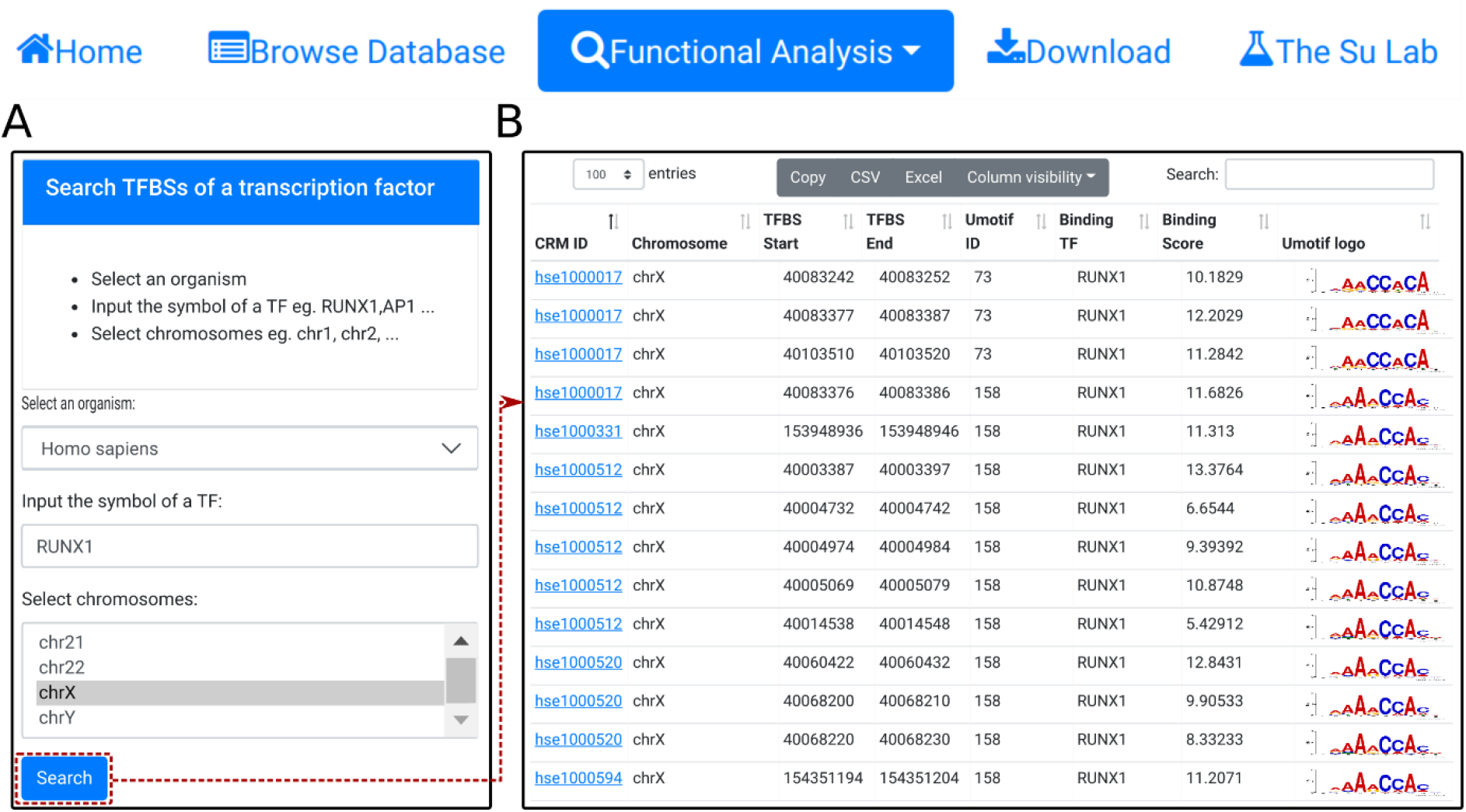
Search TFBSs of a TF. A. In the search form, the use selects an organism (e.g., *H. sapiens*), input the name of a TF (e.g., *RUNX1*), and select a chromosome (e.g., chrX). **B.** A snapshot of the resulting TFBS table containing 2678 TFBSs of RUNX1 in chrX of *H. sapiens*.

### Batch download

Using the Download function from the home page, the user can download all predicted CRMCs and constituent TFBSs in an organism in file in a BED format.

## Future development

In the future, we will add predicted CRMs and TFBSs in other important model organisms such as the worm (*C. elegans*) and the fly (*D. melanogaster*). We will also update the predictions in each organism when more data are available. We will add more information about the CRMs, including their predicted functional states (active or non-active) of the CRMs in various cell/tissue types, predicted target genes, and causal variants of complex traits and diseases by integrating more data sources.

## Conclusions

We have developed the PCRMS database that contains the most comprehensive collections of accurately predicted CRMs and constituent TFBSs in the human and mouse genome. The web interface to PCRMS allows browse, search and visualization of the CRMs and constituent TFBSs. It also provides three functional analysis modules to search the closest CRM(s) to a gene, CRM(s) in a region around a gene, and TFBSs landscape of a specific TF. The results can be inspected in interactive ways and exported in files in different formats. All the predicted CRMCs and TFBSs in an organism can be download in the BED format. PCRMS will facilitate the research community’s efforts to characterize the regulatory genomes in important organisms.

## Declarations

### Ethics approval and consent to participate

Ethics approval is not applicable to this study.

### Consent for publication

Not applicable

### Availability of data and materials

All predicted CRMCs of human and mouse can be freely downloaded at https://cci-bioinfo.uncc.edu.

### Authors’ contributions

ZS and PN conceived and designed the project. PN carried out the computational analysis and built the database and the web interface. ZS and PN wrote the manuscript. All the authors read and approved the final manuscript.

## Acknowledgements

The authors would like to acknowledge members from Office of Technical Service of College of Computing and Informatics at UNC Charlotte for security reviewing and deploying the databases.

## References

1. Davidson EH: The Regulatory Genome: Gene Regulatory Networks In Development And Evolution. Academic Press; 2006.

2. Hindorff LA, Sethupathy P, Junkins HA, Ramos EM, Mehta JP, Collins FS, Manolio TA: Potential etiologic and functional implications of genome-wide association loci for human diseases and traits. Proc Natl Acad Sci U S A 2009, 106: 9362–9367.

3. Ramos EM, Hoffman D, Junkins HA, Maglott D, Phan L, Sherry ST, Feolo M, Hindorff LA: Phenotype-Genotype Integrator (PheGenI): synthesizing genome-wide association study (GWAS) data with existing genomic resources. Eur J Hum Genet 2014, 22: 144–147.

4. Wittkopp PJ, Kalay G: Cis-regulatory elements: molecular mechanisms and evolutionary processes underlying divergence. Nat Rev Genet 2012, 13: 59–69.

5. Rubinstein M, de Souza FS: Evolution of transcriptional enhancers and animal diversity. Philos Trans R Soc Lond B Biol Sci 2013, 368: 20130017.

6. Siepel A, Arbiza L: Cis-regulatory elements and human evolution. Curr Opin Genet Dev 2014, 29: 81–89.

7. King M, Wilson A: Evolution at two levels in humans and chimpanzees. Science 1975, 188: 107–116.

8. Maurano MT, Humbert R, Rynes E, Thurman RE, Haugen E, Wang H, Reynolds AP, Sandstrom R, Qu H, Brody J, et al: Systematic Localization of Common Disease-Associated Variation in Regulatory DNA. Science 2012, 337: 1190–1195.

9. Kasowski M, Kyriazopoulou-Panagiotopoulou S, Grubert F, Zaugg JB, Kundaje A, Liu Y, Boyle AP, Zhang QC, Zakharia F, Spacek DV, et al: Extensive variation in chromatin states across humans. Science 2013, 342: 750–752.

10. Kilpinen H, Waszak SM, Gschwind AR, Raghav SK, Witwicki RM, Orioli A, Migliavacca E, Wiederkehr M, Gutierrez-Arcelus M, Panousis NI, et al: Coordinated effects of sequence variation on DNA binding, chromatin structure, and transcription. Science 2013, 342: 744–747.

11. McVicker G, van de Geijn B, Degner JF, Cain CE, Banovich NE, Raj A, Lewellen N, Myrthil M, Gilad Y, Pritchard JK: Identification of genetic variants that affect histone modifications in human cells. Science 2013, 342: 747–749.

12. Huang D, Ovcharenko I: Identifying causal regulatory SNPs in ChIP-seq enhancers. Nucleic Acids Res 2015, 43: 225–236.

13. Ward LD, Kellis M: Interpreting noncoding genetic variation in complex traits and human disease. Nat Biotechnol 2012, 30: 1095–1106.

14. Pai AA, Pritchard JK, Gilad Y: The genetic and mechanistic basis for variation in gene regulation. PLoS Genet 2015, 11: e1004857.

15. Schmidt D, Wilson MD, Spyrou C, Brown GD, Hadfield J, Odom DT: ChIP-seq: using high-throughput sequencing to discover protein–DNA interactions. Methods 2009, 48: 240–248.

16. Song L, Crawford GE: DNase-seq: a high-resolution technique for mapping active gene regulatory elements across the genome from mammalian cells. Cold Spring Harbor Protocols 2010, 2010: pdb–prot5384.

17. Buenrostro JD, Wu B, Chang HY, Greenleaf WJ: ATAC- seq: a method for assaying chromatin accessibility genome-wide. Current protocols in molecular biology 2015, 109: 21–29.

18. Simon JM, Giresi PG, Davis IJ, Lieb JD: Using formaldehyde-assisted isolation of regulatory elements (FAIRE) to isolate active regulatory DNA. Nature protocols 2012, 7: 256.

19. Schones DE, Cui K, Cuddapah S, Roh T-Y, Barski A, Wang Z, Wei G, Zhao K: Dynamic regulation of nucleosome positioning in the human genome. Cell 2008, 132: 887–898.

20. Consortium EP: The ENCODE (ENCyclopedia Of DNA Elements) Project. Science 2004, 306: 636–640.

21. Consortium EP: A user’s guide to the encyclopedia of DNA elements (ENCODE). PLoS Biol 2011, 9: e1001046.

22. Bernstein BE, Stamatoyannopoulos JA, Costello JF, Ren B, Milosavljevic A, Meissner A, Kellis M, Marra MA, Beaudet AL, Ecker JR, et al: The NIH Roadmap Epigenomics Mapping Consortium. Nat Biotechnol 2010, 28: 1045–1048.

23. Kundaje A, Meuleman W, Ernst J, Bilenky M, Yen A, Heravi-Moussavi A, Kheradpour P, Zhang Z, Wang J, Ziller MJ, et al: Integrative analysis of 111 reference human epigenomes. Nature 2015, 518: 317–330.

24. Consortium GT: The Genotype-Tissue Expression (GTEx) project. Nat Genet 2013, 45: 580–585.

25. Whitington T, Frith MC, Johnson J, Bailey TL: Inferring transcription factor complexes from ChIP-seq data. Nucleic Acids Res 2011, 39: e98.

26. Sun H, Guns T, Fierro AC, Thorrez L, Nijssen S, Marchal K: Unveiling combinatorial regulation through the combination of ChIP information and in silico cis-regulatory module detection. Nucleic Acids Res 2012, 40: e90.

27. Ha N, Polychronidou M, Lohmann I: COPS: detecting co-occurrence and spatial arrangement of transcription factor binding motifs in genome-wide datasets. PLoS One 2012, 7: e52055.

28. Rohr CO, Parra RG, Yankilevich P, Perez-Castro C: INSECT: IN-silico SEarch for Co-occurring Transcription factors. Bioinformatics 2013, 29: 2852–2858.

29. Ernst J, Kellis M: ChromHMM: automating chromatin-state discovery and characterization. Nat Methods 2012, 9: 215–216.

30. Ernst J, Kheradpour P, Mikkelsen TS, Shoresh N, Ward LD, Epstein CB, Zhang X, Wang L, Issner R, Coyne M, et al: Mapping and analysis of chromatin state dynamics in nine human cell types. Nature 2011, 473: 43–49.

31. Hoffman MM, Buske OJ, Wang J, Weng Z, Bilmes JA, Noble WS: Unsupervised pattern discovery in human chromatin structure through genomic segmentation. Nat Methods 2012, 9: 473–476.

32. Andersson R, Gebhard C, Miguel-Escalada I, Hoof I, Bornholdt J, Boyd M, Chen Y, Zhao X, Schmidl C, Suzuki T, et al: An atlas of active enhancers across human cell types and tissues. Nature 2014, 507: 455–461.

33. Khan A, Zhang X: dbSUPER: a database of super-enhancers in mouse and human genome. Nucleic Acids Res 2015.

34. Jiang Y, Qian F, Bai X, Liu Y, Wang Q, Ai B, Han X, Shi S, Zhang J, Li X, et al: SEdb: a comprehensive human super-enhancer database. Nucleic Acids Res 2019, 47: D235–d243.

35. Ashoor H, Kleftogiannis D, Radovanovic A, Bajic VB: DENdb: database of integrated human enhancers. Database (Oxford) 2015, 2015.

36. Dreos R, Ambrosini G, Cavin Perier R, Bucher P: EPD and EPDnew, high-quality promoter resources in the next-generation sequencing era. Nucleic Acids Res 2013, 41: D157–164.

37. Dimitrieva S, Bucher P: UCNEbase—a database of ultraconserved non-coding elements and genomic regulatory blocks. Nucleic acids research 2013, 41: D101–D109.

38. Visel A, Taher L, Girgis H, May D, Golonzhka O, Hoch RV, McKinsey GL, Pattabiraman K, Silberberg SN, Blow MJ, et al: A high-resolution enhancer atlas of the developing telencephalon. Cell 2013, 152: 895–908.

39. Fishilevich S, Nudel R, Rappaport N, Hadar R, Plaschkes I, Iny Stein T, Rosen N, Kohn A, Twik M, Safran M, et al: GeneHancer: genome-wide integration of enhancers and target genes in GeneCards. Database (Oxford) 2017, 2017.

40. Wang J, Dai X, Berry LD, Cogan JD, Liu Q, Shyr Y: HACER: an atlas of human active enhancers to interpret regulatory variants. Nucleic Acids Res 2019, 47: D106–d112.

41. Cai Z, Cui Y, Tan Z, Zhang G, Tan Z, Zhang X, Peng Y: RAEdb: a database of enhancers identified by high-throughput reporter assays. Database 2019, 2019.

42. Wang Z, Zhang Q, Zhang W, Lin J-R, Cai Y, Mitra J, Zhang ZD: HEDD: human enhancer disease database. Nucleic acids research 2018, 46: D113–D120.

43. Zhang G, Shi J, Zhu S, Lan Y, Xu L, Yuan H, Liao G, Liu X, Zhang Y, Xiao Y: DiseaseEnhancer: a resource of human disease-associated enhancer catalog. Nucleic acids research 2018, 46: D78–D84.

44. Wei Y, Zhang S, Shang S, Zhang B, Li S, Wang X, Wang F, Su J, Wu Q, Liu H: SEA: a super-enhancer archive. Nucleic acids research 2016, 44: D172–D179.

45. Gao T, Qian J: EnhancerAtlas 2.0: an updated resource with enhancer annotation in 586 tissue/cell types across nine species. Nucleic acids research 2020, 48: D58–D64.

46. Moore JE, Purcaro MJ, Pratt HE, Epstein CB, Shoresh N, Adrian J, Kawli T, Davis CA, Dobin A, Kaul R, et al: Expanded encyclopaedias of DNA elements in the human and mouse genomes. Nature 2020, 583: 699–710.

47. Ni P, Su Z: A map of cis-regulatory modules and constituent transcription factor binding sites in 77.5% regions of the human genome. bioRxiv 2020.

48. Visel A, Minovitsky S, Dubchak I, Pennacchio LA: VISTA Enhancer Browser--a database of tissue-specific human enhancers. Nucleic Acids Res 2007, 35: D88–92.

49. Mei S, Qin Q, Wu Q, Sun H, Zheng R, Zang C, Zhu M, Wu J, Shi X, Taing L, et al: Cistrome Data Browser: a data portal for ChIP-Seq and chromatin accessibility data in human and mouse. Nucleic Acids Res 2017, 45: D658–d662.

50. Li Y, Ni P, Zhang S, Li G, Su Z: ProSampler: an ultra-fast and accurate motif finder in large ChIP-seq datasets for combinatory motif discovery. Bioinformatics 2019.

51. Basso K, Dalla-Favera R: Roles of BCL6 in normal and transformed germinal center B cells. Immunol Rev 2012, 247: 172–183.

52. Damm F, Chesnais V, Nagata Y, Yoshida K, Scourzic L, Okuno Y, Itzykson R, Sanada M, Shiraishi Y, Gelsi-Boyer V, et al: BCOR and BCORL1 mutations in myelodysplastic syndromes and related disorders. Blood 2013, 122: 3169–3177.

